# Early impact of gestational protein restriction on nephrogenesis in male mouse offspring: Role of Autophagy and Apoptosis Mechanisms

**DOI:** 10.64898/2026.04.01.715956

**Authors:** Marina S. Folguieri, Bruno Calsa, Patrícia Aline Boer, José Antonio Rocha Gontijo

## Abstract

**Background:** Maternal protein restriction results in a 28% reduction in nephrogenic cells and nephron units in rodent offspring by the 17th day of gestation compared to adequate protein intake.

**Aims:** The present study investigates the association between growth factor expression and some developmental pathways that contribute to nephron reduction during embryonic and fetal development.

**Experimental Design:** Pregnant C57BL/6-Tg and C57BL/6J mice were assigned to either normal protein intake (NP-17%) or low protein intake (LP-6%) groups. Body weight of male offspring and kidney growth factor expression were assessed on gestation days (GD) 14 and 18.

**Results:** On GD 14, LP pups exhibited a 4% higher body mass (0.1035 g) compared to NP pups (0.0995 g, p = 0.005). By GD 18, LP pups demonstrated a 4% decrease in body mass (0.939 g, p = 0.03) and a 10% increase in the number of cells per metanephric cap area. Three genes (Csf2, Il1b, Il2) were downregulated, while seven genes (Bmp2, Csf3, Fgf8, Gdnf, Bmp7, Fgf3, Ntf3) were upregulated. By GD 14, phagophores and autophagosomes in the ureteric bud increased by 197%, with further increases observed by GD 18. Bcl-2 expression increased significantly in ureteric bud cells, and mTOR activity was elevated by GD 18.

**Conclusion:** Early gestational protein restriction modifies renal growth factor gene expression, influencing cell proliferation and autophagy, and may contribute to reduced nephron numbers by the 18th day of gestation.

**HIGHLIGHTS:**

- This study examines the effects of a low-protein diet during pregnancy in mice and demonstrates a significant reduction in embryo-fetal body weight between gestational days 14 and 18.
- Protein restriction induces a distinct cellular pattern in the mesonephros, with a 21% increase in CAP cells at gestational day 14 (GD14), followed by a decrease by gestational day 18 (GD18) compared to offspring from mothers on a normal protein diet.
- Additionally, increased expression levels of key growth factors essential for kidney development were observed at GD 14, comparing LP with NP intake during pregnancy.
- Seven genes were upregulated (Gdnf, Bmp2, Bmp7, Tgfα, Fgf8, Fgf3, Csf3, Ntf3), while three genes were downregulated (Csf2, Il1b, Il2).
- Overall, these findings indicate that gene regulation, autophagy, and mTOR signaling mechanisms significantly influence nephron numbers in response to gestational protein restriction beyond the 18th day of gestation.

## INTRODUCTION

Kidney development depends on precise genetic signals that determine nephron number and structure [1–6]. Entry of the nephric duct into the Metanephric Mesenchyme (MM) forms the Ureteral Bud (UB), which interacts with cap mesenchymal cells for nephron formation. A balance between proliferation and differentiation ensures optimal nephron count and communication, but maternal malnutrition can disrupt this, reducing nephron numbers in offspring [1,2,4–8]. Environmental influences during embryonic and fetal periods can cause low birth weight, raising future disease risk. Fetal programming shows that in *utero* conditions have lasting health effects [3,4,6,11,12], as described by Barker’s work linking low birth weight to cardiovascular risk [13,14]. The DOHaD framework explores how early development affects long-term health. Zeman’s 1967 study [15] found that protein-restricted pregnancies in rats led to fewer nephrons and smaller kidneys. Our research confirmed decreased nephron count, altered angiotensin II receptor expression, and increased glomerular tuft volume in protein-restricted offspring, linked to fewer Six2+ cells in the cap mesenchyme [1,2,4,7,8,15–18]. Cap mesenchyme stem cells may be depleted prematurely, disrupting nephron formation. Normally, some cells differentiate, and others proliferate, maintaining enough stem cells for nephron development.

Additionally, autophagy and apoptosis supply essential nutrients that support nephron growth. These mechanisms are critical during fetal development and may be disrupted by placental disorders such as intrauterine growth restriction (IUGR) [19]. The present study investigates, at two specific time points—early kidney development and near the end of gestation—the relationships among growth factors, molecular expression, and developmental mechanisms that may contribute to nephron reduction during the embryonic-fetal period in offspring of mothers with low- versus normal-protein intake throughout gestation.

## MATERIALS AND METHODS

### Animals and Ethical Procedures

The Institutional Ethics Committee approved the experimental protocol (#5481-1/2020 CEUA/UNICAMP), and the Internal Biosafety Commission authorized the use of transgenic animals (CIBIO FCM n° 08/2019). C57BL/6J and C57BL/6-Tg(CAG-RFP/EGFP/Map1lc3b)1Hill/J (CAG-RFP-EGFP-LC3) mice, aged 8 to 10 weeks, were obtained from the Multidisciplinary Center for Biological Research in Laboratory Animals at Unicamp (CEMIB). Animals were maintained at 23 ± 2°C and 50 ± 10% humidity under a 12-hour light-dark cycle. CAG-RFP-EGFP-LC3 mice express red fluorescent protein (RFP) and enhanced green fluorescent protein (EGFP), which serve as pH-sensitive markers within cellular compartments such as phagosomes. Lamp-1 (Novus Biologicals, NB120-19294, 1:2000) was used as a lysosomal marker. For the experimental model, 16-week-old female CAG-RFP-EGFP-LC3 mice were paired with male C57BL/6J mice for two hours during the dark cycle. Females were assigned to four groups based on diet and gestational age: standard (normal) protein (NP, 17% casein, n = 22) or low-protein (LP, 6% casein, n = 20), both isocaloric with standard sodium and calcium content (Table 1S). On gestational days 14 or 18 (GD), dams were anesthetized with 3% isoflurane, and only male fetuses were collected to control for sex-based differences in developmental programming. Fetuses were either fixed in 4% paraformaldehyde (PFA) and stored in 70% alcohol or frozen in N-hexane or liquid nitrogen and stored at -80°C.

### Sex Determination and Genotyping

DNA was extracted from tail fragments using 50 mM NaOH at 100°C. The presence of the SRY gene, a sex-determining gene located on the Y chromosome, was assessed by conventional PCR to determine fetal sex. The GoTaq® G2 Green Master Mix (M7823, Promega, Madison, USA) was used for 35 cycles at 95°C, 59°C, and 72°C, each for 1 minute. The SRY product (Forward: GTGAGAGGCACAAGTTGGC, Reverse: CTCTGTGTAGGATCTTCAATC) is 147 bp, while the positive control (MYG; Forward: TTACGTCCATCGTGGACAGC, Reverse: TGGGCTGGGTGTTAGTCTTA) is 246 bp.

Genotyping was necessary as CAG-RFP-EGFP-LC3 mice were crossed with wild-type C57BL/6J mice; verification of the presence of the transgene in fetuses was performed by PCR (Forward: CAG GAC GAG CTG TAC AAG T, Reverse: CAC CGT GAT CAG GTA CAA GGA for the transgene, resulting product 208 bp; oIMR7338: TTACGTCCATCGTGGACAGC, oIMR7339: TGGGCTGGGTGTTAGTCTTA as positive control, 324 bp). Polymerase chain reaction (PCR) is a technique that amplifies specific DNA sequences for identification.

### Kidney Volume Measurement

At 14 and 18 gestational days (n = 5/group), fetal kidneys were dissected and measured for volume. Fetal gestational days refer to the age of the embryo and fetus in days post-conception.

### RT2 Profiler PCR Arrays and RNA Extraction

Kidney tissues from fetuses at 14 GD (NP, n=3; LP, n=3) were processed using the RNeasy Plus Micro kit (74034, Qiagen, Hilden, Germany) to extract ribonucleic acid (RNA). RNA purity and concentration were assessed with a NanoDrop 2000c spectrophotometer. Complementary DNA (cDNA) was synthesized from 0.5 µg RNA using the RT² First Strand kit. After synthesis, 91 µL RNase-free water was added, and samples were stored at -20°C. cDNA quality was evaluated with the RT2 RNA QC PCR Array (PAMM-999ZC-1) before amplification. For gene expression analysis, RT2 Profiler PCR arrays (PAMM-041ZC, Qiagen, USA) were used for quantitative PCR (qPCR), targeting 84 growth factor-related genes (Table 3S). Amplification was performed with SYBR Green ROX qPCR Mastermix (1123342, Qiagen, USA) on a StepOne Plus thermocycler, following the protocol: denaturation at 95°C for 10 minutes, then 40 cycles of 95°C for 15 seconds and 65°C for 1 minute. Cycle threshold (Ct) values were normalized to Actb, B2m, Gapdh, and Hsp90ab1 as reference genes. This workflow outlines the process from RNA extraction to gene expression analysis.

### Immunohistochemistry

Fetuses preserved in 70% alcohol underwent dehydration, clearing, and embedding in Paraplast (Sigma-Aldrich, USA), a material used to support tissue for sectioning. Five-micrometer-thick tissue sections were further processed for immunoperoxidase or immunofluorescence, techniques that detect specific proteins using labeled antibodies. After hydration with phosphate-buffered saline (PBS, pH 7.2), which maintains pH and ionic strength, antigen retrieval was performed in a citrate buffer bath (pH 6.0) for 30 minutes to unmask epitopes for antibody binding.

For immunoperoxidase, slides were blocked with 5% non-immune serum for 1 hour, incubated overnight with the primary antibody in 3% BSA (bovine serum albumin), treated with hydrogen peroxide to inhibit endogenous peroxidase, then exposed to a secondary antibody and DAB (3,3’-diaminobenzidine) to visualize the antigen (brown coloration). Slides were dehydrated and mounted with Entellan®. For immunofluorescence, slides were blocked with 5% non-immune serum and 0.2% Triton X-100 in PBS, incubated overnight with the primary antibody, washed, and incubated with the secondary antibody (Table 2S). DAPI (4′,6-diamidino-2-phenylindole, 0.1 µg/mL) was included in the final wash to stain nuclei.

All images were acquired using an Airyscan Inverted Microscope at INFABIC, State University of Campinas, and analyzed with ImageJ, a software program for image processing. Negative controls, which omit the primary antibody to assess specificity, showed no immunoreactivity.

### Quantification

Tissue sections were analyzed to quantify the nephrogenic area, cap mesenchyme (CM), and ureteral bud (UB) in embryos at 14 and 18 gestational days (GD) from low-protein (LP, n = 4) and normoprotein (NP, n = 4) groups, each derived from different mothers. All CM and UB structures were quantified in four NP and four LP metanephros samples.

### Autophagic Flux

Images of CM and UB were acquired using an EC Plan-Neofluar 40x/1.3 Oil DIC objective on an Airyscan confocal microscope (Carl Zeiss) with 1.8x zoom. The microscope was equipped with laser lines for DAPI (405 nm), eGFP (488 nm), mRFP (543 nm), and Alexa 647 to optimize imaging for quantification of autophagic vesicles. ImageJ software was used to automate vesicle type and count analysis through a defined command sequence. Mean intensity across red, green, and blue channels was measured and classified by HUE angle (Yellow, Orange, Red, Purple, Blue). Vesicle counts were normalized to the CM and UB area (vesicles per mm²).

### Data Presentation and Statistical Analysis

All quantitative data are mean ± standard deviation (SD). Temporal analyses used one-way ANOVA with Bonferroni’s post hoc test. Two-group comparisons used two-way repeated-measures ANOVA and Tukey’s post hoc test. Student’s t-tests compared independent groups, with Welch’s t-test for heteroscedasticity. Survival analysis used Mantel-Cox and Gehan-Breslow-Wilcoxon tests. Analyses were done with GraphPad Prism 5.00 (GraphPad, USA); p ≤ 0.05 was significant.

## RESULTS

### Maternal Parameters

Table 1 presents maternal protein restriction (LP offspring) during the 14th and 18th gestational days (GD) compared to the normal protein (NP) group. At 14 GD, LP offspring showed a 24% decrease in weight gain and caloric efficiency. By 18 GD, the LP group showed a 32% greater decrease in weight gain and food efficiency, with a 13% increase in feed consumption (Table 1).

**Table 1.** Maternal Parameters with 14 and 18 gestational days (GD).

| Gestational age | Group | Weight gain (g) |  | Feed consumption (g) |  | Feed efficiency |  |
| --- | --- | --- | --- | --- | --- | --- | --- |
| | | Mean ± SD (n) | p value | Mean ± SD | p value | Mean ± SD | p value |
| 14<sup>th</sup> GD | NP | 9,23 ± 0,9 (11) | <0,0001 | 47,19 ± 2,7 | 0,125 | 9,23 ± 0,9 | 0,0063 |
| | LP | 6,97 ± 0,7 (9) | | 48,63 ± 2,5 | | 6,97 ± 0,7 | |
| 18<sup>th</sup> GD | NP | 16,15 ± 1,7 (11) | <0,0001 | 59,07 ± 8,6 | 0,019 | 0,067 ± 0,17 | 0,039 |
| | LP | 10,89 ± 1,4 (11) | | 67,13 ± 6,4 | | 0,044 ± 0,01 | |

### Fetal Parameters

Maternal protein restriction during gestation led to significant body weight differences between male and female fetuses. Placental mass and the fetus-to-placental ratio were unchanged in both groups. At 14 GD, LP male and female fetuses showed body mass increases of 6% and 9%, respectively, compared with NP progeny. At 18 GD, body mass decreased by 4% in male and 8% in female LP fetuses (Table 2).

**Table 2.** The effects of maternal protein restriction during gestation on fetal parameters.

| Gestational age | Sex | Group | Fetal body mass (g) |  | Placental mass (g) |  | f:p ratio (g/g) |  |
| --- | --- | --- | --- | --- | --- | --- | --- | --- |
| | | | Mean ± SD (n) | p value | Mean ± SD | p value | Mean ± SD | p value |
| 14<sup>th</sup> GD | Male | NP | 0,199 ± 0,02 (46) | 0,005 | 0,103 ± 0,03 | 0,5 | 2,074 ± 0,67 | 0,07 |
| | | LP | 0,211 ± 0,02 (40) | | 0,099 ± 0,02 | | 2,286 ± 0,66 | |
| | Female | NP | 0,189 ± 0,02 (41) | 0,001 | 0,097 ± 0,03 | 0,7 | 2,049 ± 0,62 | 0,2 |
| | | LP | 0,206 ± 0,02 (35) | | 0,099 ± 0,03 | | 2,163 ± 0,59 | |
| 18<sup>th</sup> GD | Male | NP | 0,979 ± 0,08 (49) | 0,03 | 0,115 ± 0,03 | 0,9 | 8,983 ± 2,3 | 0,2 |
| | | LP | 0,939 ± 0,09 (39) | | 0,116 ± 0,02 | | 8,411 ± 1,5 | |
| | Female | NP | 0,977 ± 0,89 (27) | 0,0006 | 0,111 ± 0,03 | 0,09 | 8,917 ± 2,1 | 0,2 |
| | | LP | 0,895 ± 0,08 (44) | | 0,099 ± 0,03 | | 9,520 ± 2,2 | |

### Kidney Development

Analysis focused on male offspring (see Discussion). No significant kidney volume changes occurred at either gestational age (Figure 1). At 14 GD, LP fetuses’ metanephros had a 21% higher Cap mesenchymal (CAP) cell count than NP. At 18 GD, the CAP cell count was similar between groups (Figure 1).

**Figure 1.**
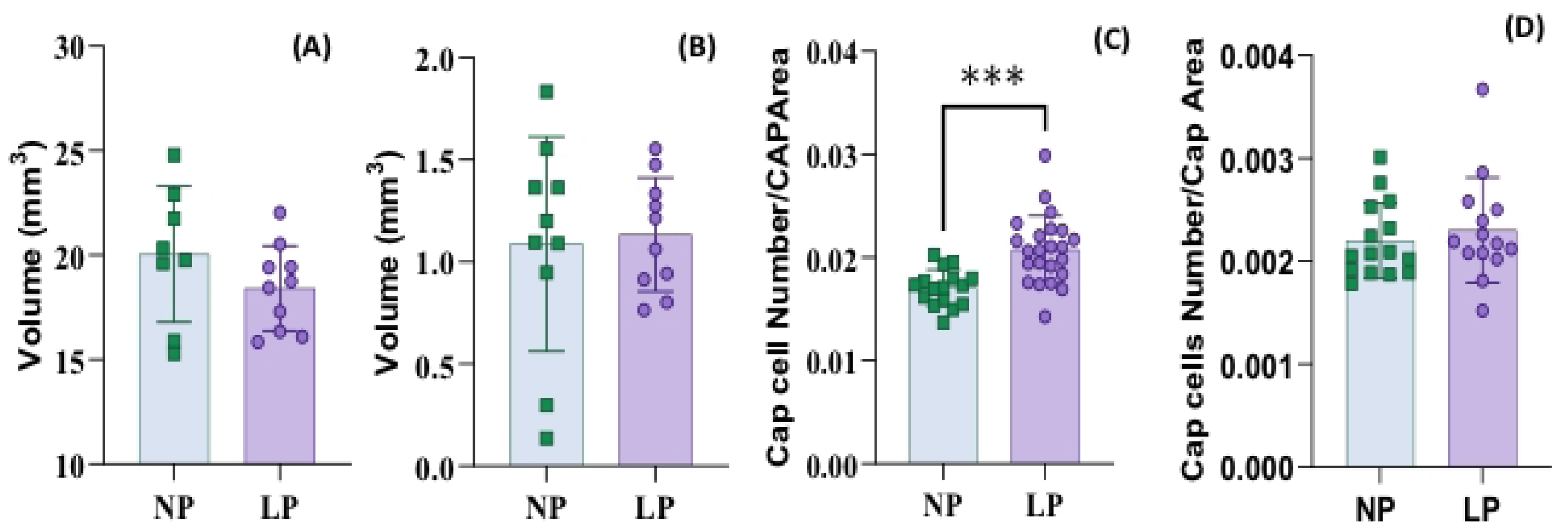
shows kidney volume at (A) 14 gestational days (GD), where the NP group measured 1.089 ± 0.5259 (n = 10) and the LP group measured 1.134 ± 0.278 (n = 10), with a p-value of 0.8128. At (B) 18 GD, the NP group’s volume was 20.04 ± 3.249 (n = 8) compared to the LP group’s 18.41 ± 2.032 (n = 10), resulting in a p-value of 0.2119. For the number of cells in the Cap Area: © at 14 GD, the NP offspring had 0.01705 ± 0.001773 cells (n = 16), and the LP offspring had 0.02076 ± 0.003311 cells (n = 23), with a p-value of 0.0002. At (D) 18 GD, the NP group had 0.002203 ± 0.0003669 cells (n = 15), and the LP group had 0.002304 ± 0.0005117 cells (n = 14), with a p-value of 0.5453.

### PCR Array of Growth Factors

PCR array analysis was performed on the kidneys of male offspring at 14 GD, covering 84 genes. Ten genes were differentially expressed (Table 2S). Two genes (Csf2, Il2) were downregulated, and eight (Bmp2, Csf3, Fgf8, Gdnf, Bmp7, Fgf3, Ntf3, Tgfα) were upregulated in LP compared to NP offspring (Figures 2 and Table 3S).

**Figure 2.**
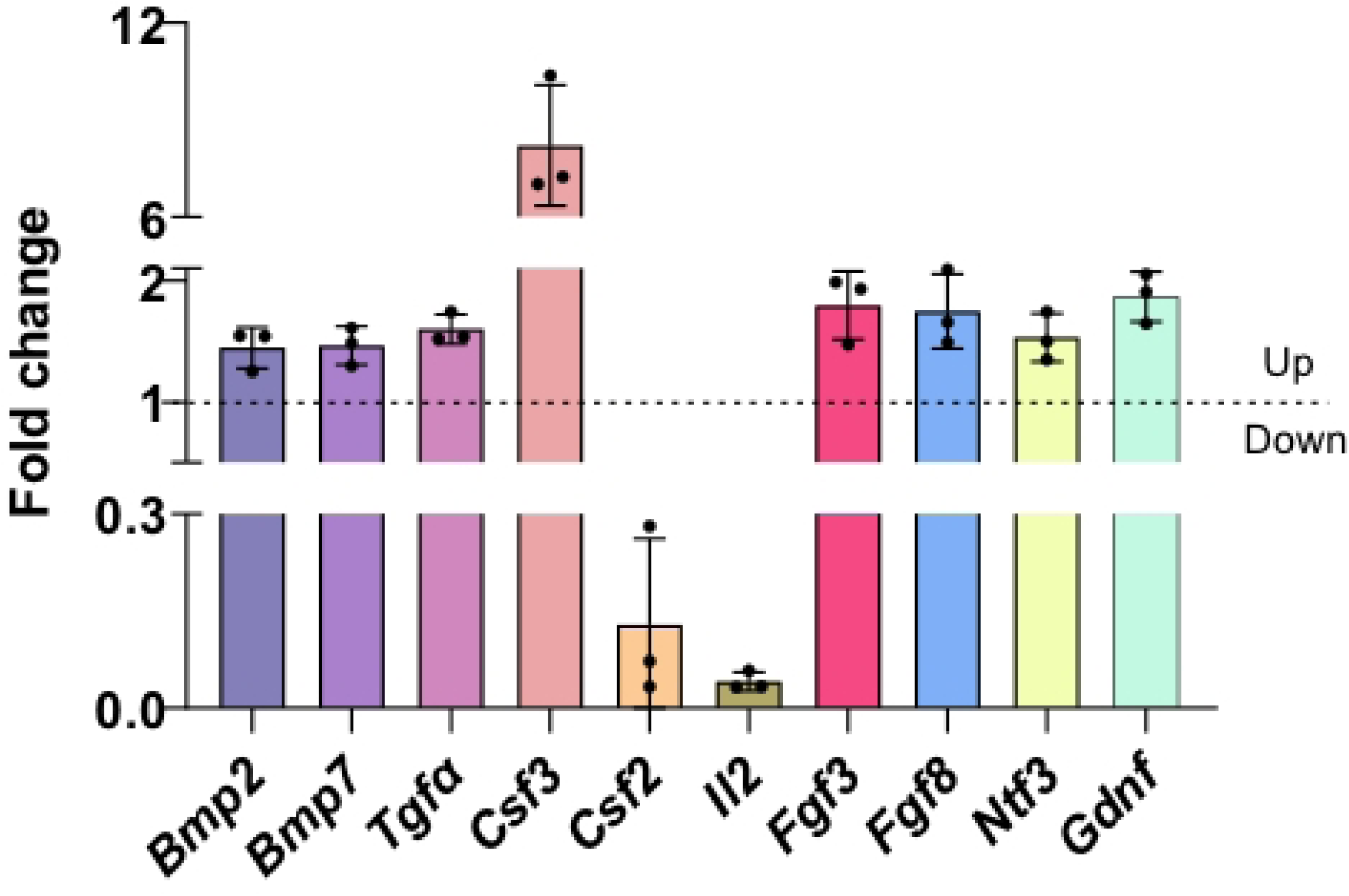
Growth factors are differentially expressed in the kidney at 14 GD. Three genes were downregulated (Csf2, Il1b, Il2), and seven genes were upregulated (Bmp2, Csf3, Fgf8, Gdnf, Bmp7, Fgf3, Ntf3).

### Autophagy Assessment

Autophagy in the metanephros was evaluated by HUE scale color classification (Figure 3). At 14 GD, phagophore and autophagosome (yellow vesicle) numbers in UB cells rose by 197%, and lysosomes (blue) in CAP cells by 270% in LP versus NP fetuses (Table 3, Figure 3). At 18 GD, LP UB cells had a 246% rise in phagophores/autophagosomes, a 92% decrease in mature autophagosomes (red/purple), an 82% drop in autolysosomes, and a 119% increase in CAP vesicle count compared to NP (Table 2S, Figure 3).

**Figure 3.**
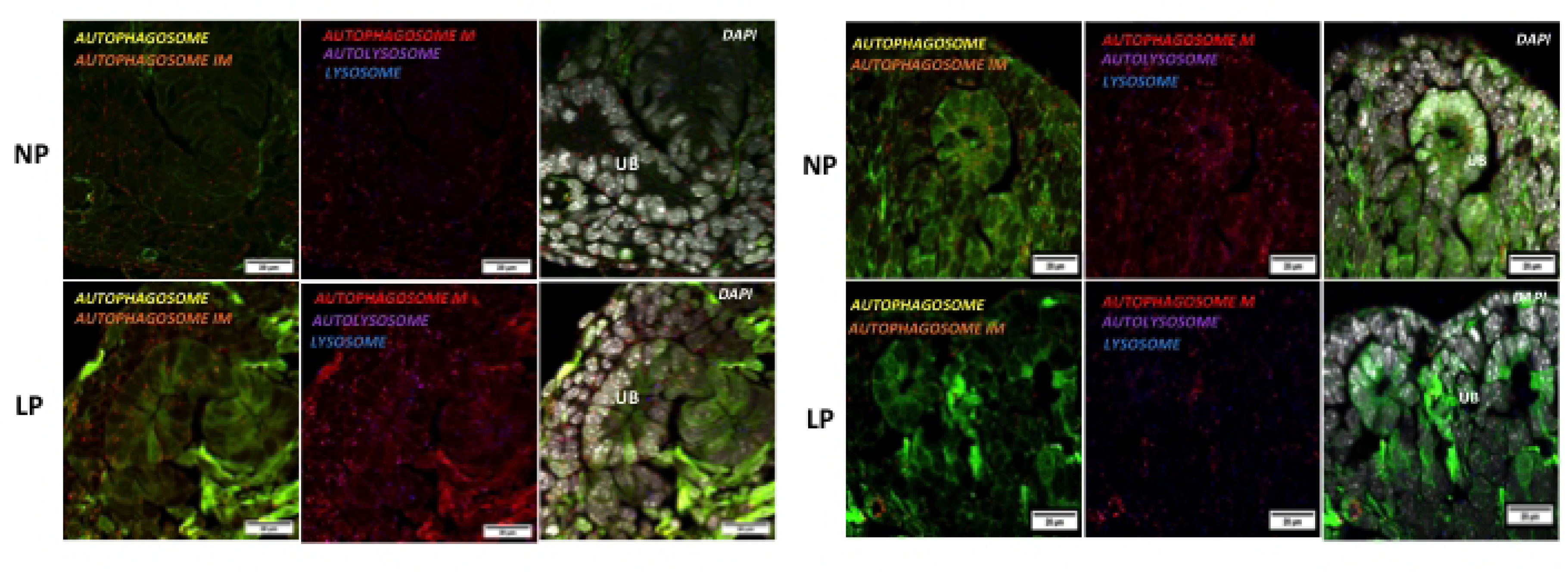
illustrates the autophagy process at 14 gestational days (GD) in both the normal protein (NP) and low protein (LP) intake offspring groups. At 14 GD, there was a 197% increase in the number of autophagosomes in the ureteric bud (UB) cells, along with a 270% increase in lysosomes in CAP cells. The autophagy process was also observed at 18 GD in both the NP and LP offspring. During this later stage, there was an increase in the number of autophagosome vesicles; however, there was a decrease in the maturation of autophagosomes (M) and a reduction in the number of autolysosomes in the ureteric bud.

**Table 3.** Autophagy process with 14 GD and 18 GD.

| Gestational age | Location | Group | Autophagosome |  | Autolysosome |  | Autolysosome |  | Autolysosome with LAMP 1 |  | Lysosome |  |
| --- | --- | --- | --- | --- | --- | --- | --- | --- | --- | --- | --- | --- |
|  |  |  | Mean ± SD | p value | Mean ± SD | p value | Mean ± SD | p value | Mean ± SD | p value | Mean ± SD | p value |
| 14<sup>th</sup> GD | BU | NP | 0.74±0.5 | 0.004* | 0.56±1 | 0.9 | 0.55±0.25 | 0.2 | 1.12±0.42 | 0.4 | 6.52±6.22 | 0.3 |
|  |  | LP | 2.21±0.9 |  | 1.68±1 |  | 0.40±0.19 |  | 1.59±1.4 |  | 3.93±3.6 |  |
|  | cap | NP | 1.79±1.7 | 0.5 | 1.74±1.5 | 0.2 | 0.88±0.31 | 0.7 | 1.98±1.3 | 0.7 | 0.997±0.2 | 0.02* |
|  |  | LP | 1.21±1.1 |  | 0.97±0.7 |  | 1.02±0.91 |  | 1.78±1.2 |  | 3.714±2.4 |  |
| 18<sup>th</sup> GD | BU | NP | 14.48±17.0 | 0.04* | 3.38±3.1 | 0.1 | 0.26±0.24 | 0.02* | 0.57±0.5 | 0.03* | 1.86±2.3 | 0.07 |
|  |  | LP | 50.21±42.8 |  | 7.013±4.2 |  | 0.02±0.04 |  | 0.1±0.1 |  | 0.26±0.32 |  |
|  | cap | NP | 11.87±8.57 | 0.1 | 6.79±6.8 | 0.08 | 0.36±0.26 | 0.2 | 0.56±0.4 | 0.27 | 1.97±2.7 | 0.3 |
|  |  | LP | 27.15±24 |  | 17.01±12.3 |  | 0.72±0.75 |  | 0.38±0.2 |  | 0.85±0.57 |  |

### Autophagy Controllers

At 14 GD, mTOR and AMPKα immunoreactivity did not differ between LP and NP metanephros (Figure 4). At 18 GD, mTOR reactivity increased by 43% in UB and 158% in CAP cells in LP metanephric regions. AMPKα activity also increased by 58% in UB and 76% in CAP cells in LP (Figure 4).

**Figure 4.**
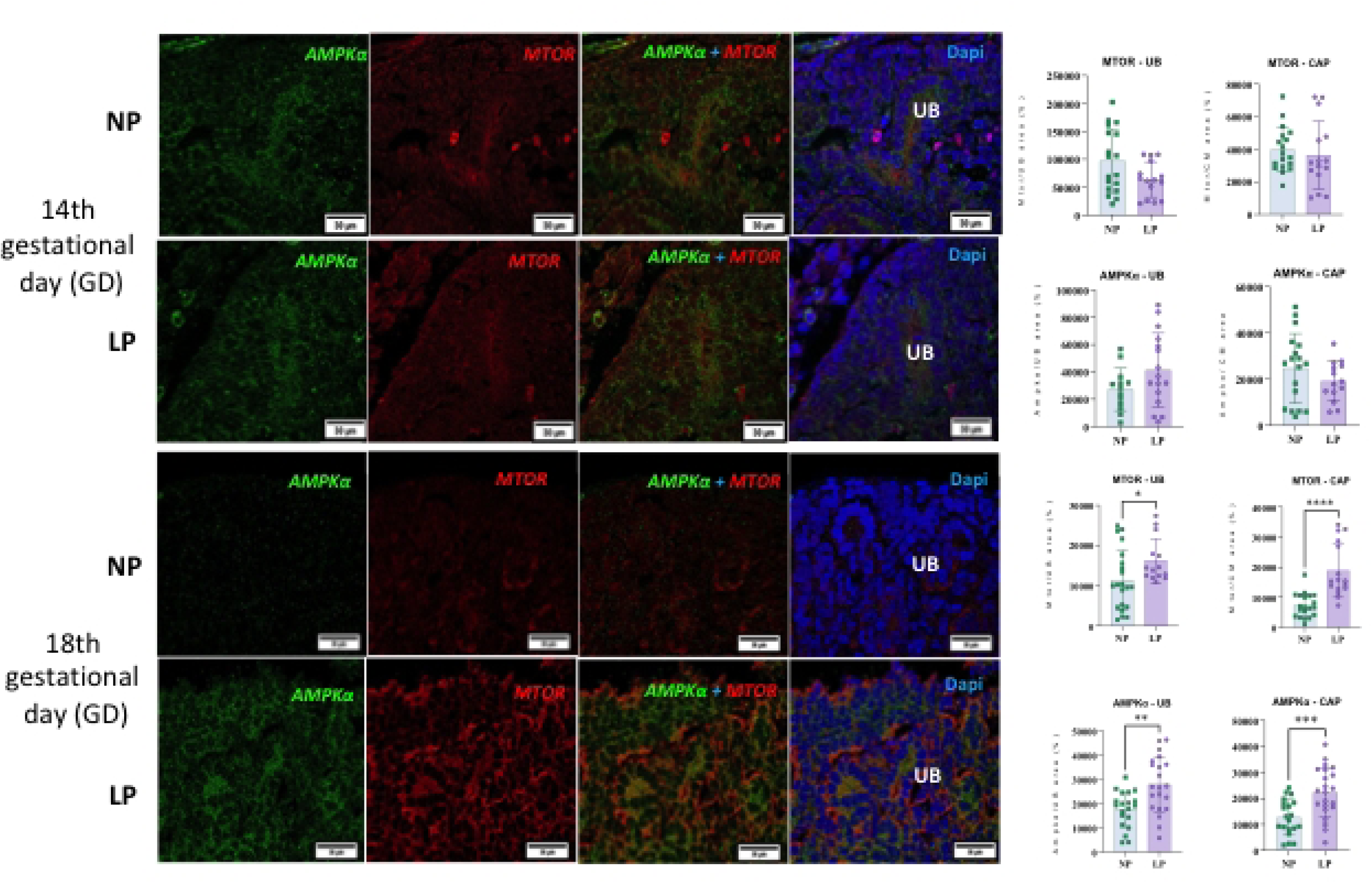
presents the analysis of mTOR and AMPKα, along with merged images of mTOR, AMPKα, and DAPI in the ureteric bud (UB) and the Cap at 14 gestational days. The results for mTOR in the UB are as follows: Normal Protein (NP) group: 96814 ± 56862, n = 18; Low-Protein (LP) intake group: 63715 ± 31661, n = 15; p = 0.0536. For mTOR in the Cap: NP group: 39682 ± 13742, n = 18; LP group: 36378 ± 20757, n = 15; p = 0.5881. In the case of AMPKα in the UB: NP group: 27243 ± 15502, n = 13; LP group: 41270 ± 27102, n = 16; p = 0.1094. For AMPKα in the Cap: NP group: 24736 ± 15098, n = 14; LP group: 19155 ± 8592, n = 18; p = 0.2269. Moving on to the analysis at 18 gestational days, the results for mTOR in the UB are: NP group: 11358 ± 7515, n = 21; LP group: 16254 ± 5460, n = 14; p = 0.0441. For mTOR in the Cap: NP group: 7306 ± 3953, n = 20; LP group: 18874 ± 8875, n = 16; p < 0.0001. For AMPKα in the UB: NP group: 17512 ± 7444, n = 20; LP group: 27785 ± 11478, n = 21; p = 0.0017. Finally, for AMPKα in the Cap: NP group: 12659 ± 6811, n = 20; LP group: 22310 ± 9589, n = 22; p = 0.0006.

### Apoptosis and Proliferation

At 14 GD, activated Caspase 3 fell by 50% in LP UB (Figure 5), remaining unchanged at 18 GD versus NP. PCNA showed no metanephron change at 14 GD, but at 18 GD, levels decreased by 55% in UB and 51% in CAP of LP fetuses (Figure 5). Bcl-2 expression rose by 2897% in UB and 443% in CAP in LP fetuses, with no change at 18 GD compared to NP (Figure 6).

**Figure 5.**
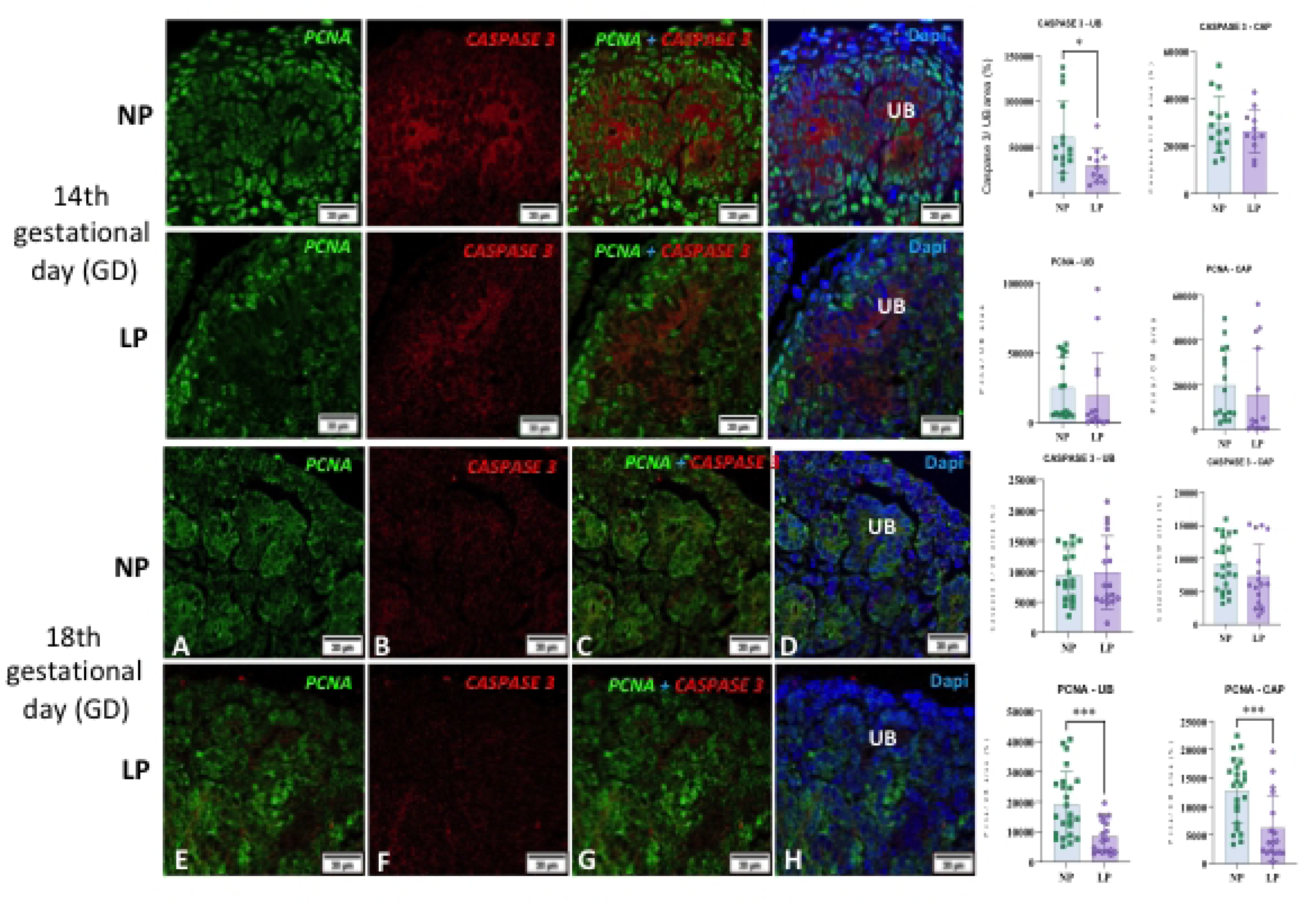
Analysis of caspase 3, PCNA, and the merged images of caspase 3, PCNA, and DAPI in the ureteric bud (UB) and Cap (CAP) at 14 gestational days. Caspase 3 UB: NP, 61112±39327, n=15; LP, 30105±18947, n=11; p= 0.0241. Caspase 3 CAP: NP, 29204±11841, n=15; LP, 26225±9165, n=11; p= 0.4940. PCNA UB: NP, 25162±21295, n=16; LP - 19528±30476, n=14; p= 0.5580. PCNA CAP: NP, 19811±15901, n=16; LP, 15573±20541, n=14; p= 0.5298. Caspase and PCNA were merged using Caspase-3, PCNA, and DAPI in UB and Cap at 18 gestational days. For Caspase-3 in the UB: NP offspring exhibited a mean of 9291±4267 (n=19), and the LP offspring showed 9823±5956 (n=17) with a p-value of 0.7581. For Caspase in the first plate, the NP intake group had 9160±3962 (n=23), and the LP intake group had 7234±4854 (n=17) with a p-value of 0.1753. For PCNA in the UB, the NP group scored 19009 ± 10917 (n = 24), while the LP group scored 8547 ± 5555 (n = 18), with a p-value of 0.0006. For PCNA in the CAP, the NP had 12724 ± 5680 (n = 23), and the LP offspring showed 6181 ± 5792 (n = 17), with a p-value of 0.0010.

**Figure 6.**
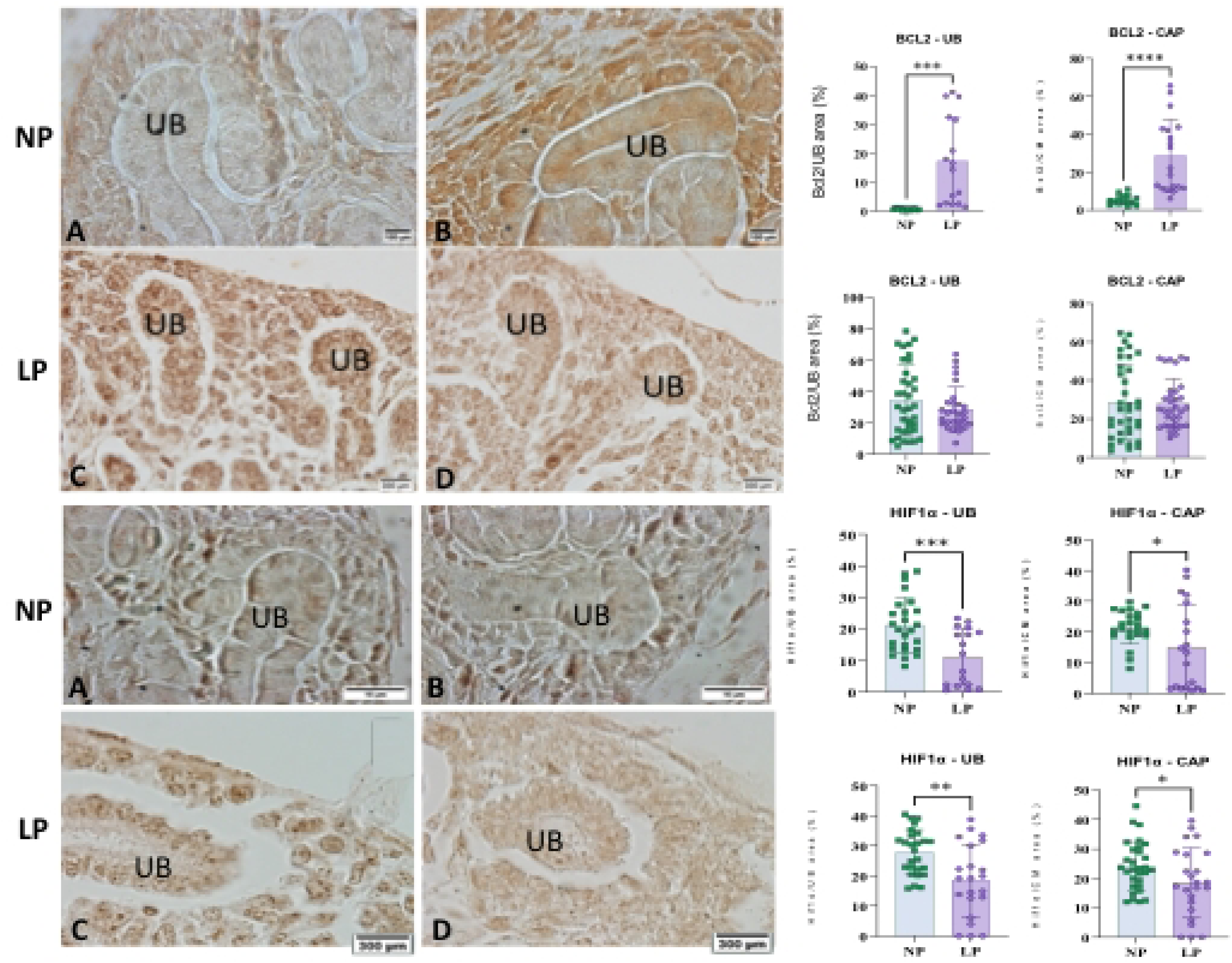
Bcl-2 levels in Cap and UB at 14 gestational days (A and B): (A) NP and (B) LP offspring. In the UB, the NP group showed a mean of 0.5735 ± 0.347 (n = 13), and the LP intake group showed a mean of 17.19 ± 15.13 (n = 16), with a p-value of 0.0005. In the CAP, the NP intake group had a mean of 5.279±2.554 (n=15), while the LP group reported a mean of 28.71±19.20 (n=19), with a p-value of <0.0001. For Bcl-2 levels in Cap and UB at 18 gestational days (C and D): © NP group and (D) LP group. In the UB, the NP group had a mean of 34.20±22.46 (n=36), and the LP group showed a mean of 28.62±14.19 (n=29), with a p-value of 0.2494. In the CAP, the NP group scored 28.68±19.40 (n=36), and the LP group had 27.97±12.94 (n=33) with a p-value of 0.8606. Displays HIF1α levels in the Cap and UB tissues at 14 gestational days, shown in panels A and B for the NP and LP offspring. For the UB tissue, the NP group showed HIF1α levels of 21.13 ± 8.704 (n = 25), while the LP group had levels of 11.07 ± 8.918 (n = 17), with a p-value of 0.0008. In the Cap tissue, the NP group recorded levels of 21.43 ± 5.466 (n = 19), while the LP group had levels of 14.92 ± 13.99 (n = 19), with a p-value of 0.0465. HIF1α levels in Cap and UB tissues at 18 gestational days are shown in panels C and D: panel C for the NP and panel D for the LP offspring. For the UB tissue at this point, the NP group had HIF1α levels of 27.9 ± 7.489 (n = 26), while the LP group had levels of 18.5 ± 11.95 (n = 22), with a p-value of 0.0018. In the Cap tissue, the NP group reported levels of 24.12 ± 8.341 (n = 30), while the LP group had levels of 18.6 ± 11.76 (n = 24), with a p-value of 0.0489.

### Immunoreactivity of hypoxia-inducible factor HIF-1α

The immunoreactivity of hypoxia-inducible factor HIF-1α was significantly diminished in UB (47%) and CAP (30%) cells of the metanephron from LP fetuses at 14 GD (Figure 6). At 18 GD, HIF-1α expression remained lower in UB (33%) and CAP (22%) cells (Figure 6) compared to the NP group. Nonetheless, the vascular endothelial growth factor (VEGF) induced by HIF-1α demonstrated a substantial increase at 14 GD in UB (458%) and CAP (120%) cells, remaining unchanged at 18 GD in LP compared to NP fetuses (Figure 7).

**Figure 7.**
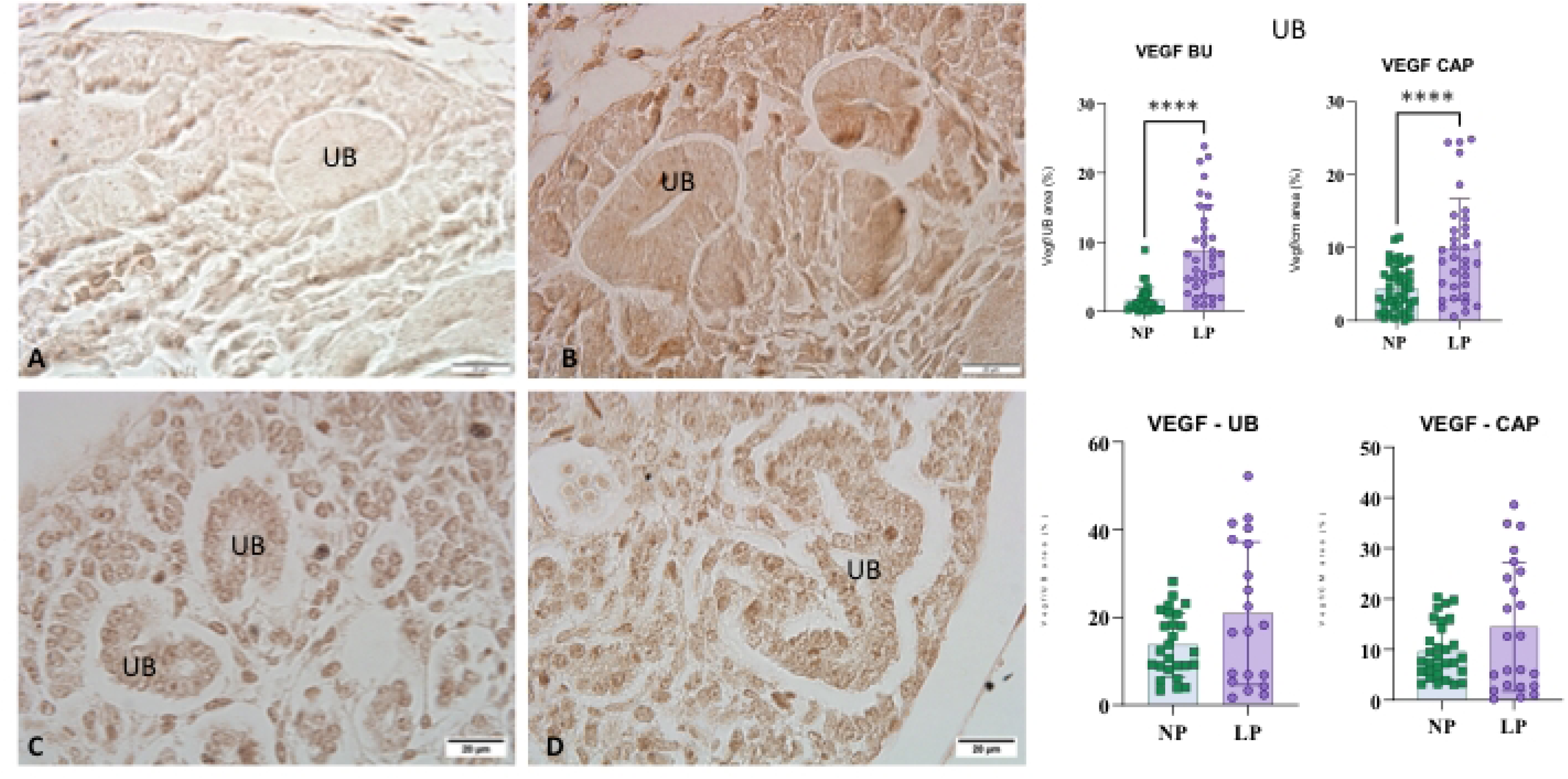
Illustrates the levels of VEGF in the Cap and UB tissues at 14 gestational days, shown in panels A and B for the NP and LP intake offspring, respectively. For the UB tissue, the NP group had VEGF levels of 1.587 ± 1.848 (n = 23), while the LP group showed levels of 8.836 ± 6.507 (n = 36), with a p-value of < 0.0001. In the Cap tissue, the NP group recorded levels of 4.44 ± 3.08 (n = 32), while the LP group had levels of 9.763 ± 6.911 (n = 36), with a p-value of < 0.0001. At 18 gestational days, VEGF levels in the Cap and UB tissues are presented in panels C and D, with panel C representing the NP group and panel D representing the LP group. For the UB tissue at this point, the NP group had VEGF levels of 13.74 ± 7.254 (n = 26), whereas the LP group had levels of 20.92 ± 16.32 (n = 20), with a p-value of 0.0511. In the Cap tissue, the NP group reported 9.623 ± 5.449 (n = 30), while the LP group reported 14.45 ± 12.73 (n = 23), with a p-value of 0.0674.

## DISCUSSION

The impact of a low-protein diet during pregnancy in mice was assessed, demonstrating a significant reduction in embryo-fetal body weight between gestational days 14 and 18. Female mice in the low-protein (LP) group exhibited increased food intake on day 18 of gestation; however, this did not improve caloric efficiency or weight gain compared to the control group [20–23]. Protein-rich diets typically enhance satiety and reduce total caloric intake, whereas low-protein diets often lead to increased food intake. These findings align with previous studies on weight gain and food intake in pregnant populations [24–30]. Gestational protein restriction induced a distinct cellular mesonephric pattern, associated with changes in fetal body weight. Specifically, a 21% increase in CAP cells was observed at 14 gestational days (GD), followed by a decrease at 18 GD, compared with offspring from a normal protein (NP) diet. This reversal in LP offspring was accompanied by decreased body weight and no significant difference in CAP cells compared with the NP group. These results are consistent with established associations between gestational protein restriction, intrauterine growth restriction, and low birth weight [1,2,8,24,31]. Previous research suggests that protein restriction may initially increase fetal body weight, but this trend reverses as gestation progresses, resulting in decreased fetal weight in protein-restricted groups [1,2,8,16,17,32]. The adaptive nature of maternal and fetal circulations is evident, as placental efficiency adjusts in response to protein restriction. Some models propose that protein restriction may enhance placental efficiency to support fetal growth early in gestation. Still, nutrient availability may become insufficient as gestation advances, leading to intrauterine growth restriction [33].

Placenta-fetal phenotypic plasticity may account for the increased fetal weight and higher CAP cell numbers observed at 14 days of gestation (GD). The placenta responds to protein restriction by upregulating amino acid and glucose transporters to support fetal development. However, inadequate nutrient supply can result in intrauterine growth restriction, as demonstrated at 18 days of gestation. No significant differences in the kidney-to-body weight ratio were detected at the gestational ages examined, in contrast to findings in newborn or older rodents subjected to protein restriction [34–37]. The small size of fetal kidneys limited the feasibility of conventional weighing methods, potentially contributing to discrepancies with previous studies. Elevated expression of key growth factors involved in differentiation, proliferation, and branching morphogenesis was detected in embryonic and fetal kidneys at 14 gestational days in the LP group compared to the NP progeny. Seven genes were upregulated (Gdnf, Bmp2, Bmp7, Tgfα, Fgf8, Fgf3, Csf3, Ntf3), while three genes were downregulated (Csf2, Il1b, Il2). Glial cell line-derived neurotrophic factor (Gdnf) is a major inducer of ureteric bud formation, promoting growth, branching, and differentiation of CAP cells [38–40]. The Bone Morphogenetic Protein (BMP) family, particularly BMP7, is essential for kidney development by supporting the survival and proliferation of metanephric mesenchyme cells.

In contrast, Bmp2 is expressed in peritubular aggregates and early tubules, facilitating cellular differentiation [40–45]. Transforming Growth Factor Alpha (Tgfα) also showed increased expression levels associated with branching morphogenesis in the LP group. This factor is essential for branching the ureteric bud and differentiating progenitor cells into nephron structures, such as glomeruli and tubules [46]. Fibroblast growth factors (FGFs) are also significant in nephrogenesis, regulating critical processes like cell survival, migration, and differentiation [47–50]. For instance, Fgf8, expressed in the metanephric mesenchyme, has been linked to decreased kidney size and nephron formation in knockout mice [49,51]. In contrast, Fgf3 knockout mice do not exhibit significant kidney defects, suggesting that FGF gene expression may not be crucial for nephrogenesis or that other growth factors can compensate for their absence [50,51]. Neurotrophin-3 (Ntf3) has not been specifically assigned a role in renal development, although its expression is primarily observed in differentiating collecting tubules and continues throughout their maturation. Unlike other factors involved in the early stages of nephrogenesis, Ntf3 is associated with later stages of nephron organization [52].

Additionally, gene-expressed factors such as Csf2, Csf3, and Il2 have less defined roles in kidney development. These cytokines regulate the production and function of granulocytes and macrophages. Csf3 is critical for granulocyte function and contributes to glomerular mesangial cell formation [53,54]. Research suggests macrophages support kidney vasculature assembly, enhancing branching morphogenesis and nephrogenesis. The prominence of Csf3 expression in LP progeny studies indicates that Csf2 and Il2, which are involved in T-cell proliferation, may be less critical in this context [55–58].

The observed increase in growth factors that promote cell proliferation, differentiation, survival, and kidney structure maintenance raises questions regarding the presence of signaling or defense mechanisms that may trigger rapid growth in response to nutrient deficiency. Importantly, no significant increase in the cell proliferation marker PCNA was detected at 14 days of gestation; instead, a marked decrease in renal cells was observed at both 14 and 18 days of gestation. This pattern suggests that elevated growth factor levels primarily enhance the differentiation and survival of CAP cells rather than stimulating proliferation. By day 18, additional constraints may have further limited the proliferative capacity of renal cells. Previous studies have associated reduced nephron numbers resulting from intrauterine growth restriction with increased autophagy and apoptosis [59–61]. Numerous investigations of autophagy utilize markers such as LC3, which indicate the presence of autophagosomes or proteins that facilitate autophagy. In this study, transgenic animals with CAG promoter/enhancer sequences were used to express pH-dependent fluorescent proteins in conjunction with the microtubule-associated protein 1 light chain 3 alpha (Map1lc3a, or LC3) gene, which labels phagosomes. The combination of green fluorescent protein (GFP) and red fluorescent protein (RFP) enables tracking of autophagy progression, producing a yellow signal at high pH within phagophores and immature autophagosomes.

Figure 4 illustrates that the acidification of autophagosomes begins before their fusion with lysosomes, as indicated by an orange fluorescence signal [62,63]. During maturation, the signal from Enhanced Green Fluorescent Protein (EGFP) decreases, leading to the exclusive emission of a RFP signal from phagophores and autophagosomes. This process results in the formation of autolysosomes by lysosomal fusion. Notably, about 14% of the EGFP remaining in the autolysosome, characterized by a pH of 4-5, still emits weak green fluorescence [64]. The visualization used the lysosomal marker Lamp-1. Following fusion with lysosomes, autolysosomes exhibit purple fluorescence, which facilitates tracking of autophagy progression. At both 14 and 18 days of gestation, an increase in autophagosomes was observed during ureteric bud (UB) formation, indicating autophagy initiation in both experimental groups. A significant increase in lysosomes was noted solely in CAP cells. By 18 days of gestation in LP progeny, there was a marked decline in mature autophagosomes and autolysosomes, suggesting potential impairments in acidification and the signaling pathways regulating autophagy. Furthermore, this study identified key regulators of the autophagy pathway, including AMP-activated protein kinase (AMPK), which promotes the formation of autophagosomes and helps cells manage low energy levels.

The mammalian target of rapamycin (mTOR) inhibits the formation of new autophagosomes, and its increased activity at 18 days of gestation suggests a disruption in the final phase of autophagy. This may hinder the fusion of autophagosomes with lysosomes in LP offspring compared to NP nephrogenesis. Current research indicates that mTOR may obstruct the assembly of the necessary fusion complex and limit protein recruitment to the autophagosome membrane in LP animals at this stage. Consequently, this could compromise vesicle maturation. Indicates that mTOR may obstruct the assembly of the necessary fusion complex and limit protein recruitment to the autophagosome membrane in LP animals at this stage. Consequently, this could compromise vesicle maturation [65,66]. Notably, no significant differences in regulatory protein levels were observed at 14 gestational days. The present study found elevated Bcl-2 levels in BU and CAP at 14 days of gestation in LP progeny, correlating with decreased caspase-3 activation.

The Bcl-2 protein is essential for inhibiting apoptosis and may impede autophagy progression by binding to Beclin-1, potentially disrupting autophagosome formation in low-protein progeny. Despite elevated Bcl-2 levels, the regulatory complex that facilitates the fusion of autophagosomes and lysosomes may still function effectively. Additionally, the observed decrease in caspase-3 activation in 14 GD LP offspring is associated with the disruption of autophagy, primarily due to increased Bcl-2 levels [67–70]. As cellular proliferation accelerates and vascular development occurs, imbalances between oxygen supply and demand, particularly in certain regions of the developing kidney, may lead to relative hypoxia [71]. This condition can activate Hypoxia-Inducible Factor (HIF), which is critical for managing hypoxic responses and regulating genes important for energy metabolism, angiogenesis, cell proliferation, and apoptosis. HIF-1α is a key factor in tubulogenesis, as shown by its expression in the collecting ducts of both the medullary and cortical regions, as well as in glomerular cells [72–74]. Gestational protein restriction may temporarily increase maternal-to-fetal oxygen flow, thereby reducing areas with low oxygen concentrations. This phenomenon may hinder the activation of the HIF-1α pathway, which is reduced in LP offspring at 14 gestational days. Previous research highlights the critical role of HIF-1α in nephrogenesis within fetal programming models.

The reduction in cellular proliferation and rapid cell differentiation at this stage may render HIF-1α less relevant in LP fetuses by 18 GD. Although no statistically significant increase in HIF-1α levels was detected in progeny at 18 gestational days (GD), the elevation in vascular endothelial growth factor (VEGF) levels at 14 GD may be associated with increased neutrophil populations stimulated by higher colony-stimulating factor 3 (CSF3) levels. This process may enhance TGF-α levels, initiating a signaling cascade that promotes VEGF expression, improves angiogenesis, and supports the survival of renal progenitor cells [71,75,76]. Although the placenta was not directly analyzed in this study, maternal-placental-fetal phenotypic plasticity may account for the observed changes in placental efficiency during early pregnancy. Cell proliferation occurred before 14 days of gestation, as indicated by increased fetal weight and metanephric cell count at GD 14. As proliferation ceases, growth factors become critical for cell differentiation, survival, and the inhibition of cell death and autophagy via BCL2. Placental efficiency may decline when nutrient intake is insufficient to meet fetal growth demands, rendering the fetus susceptible to the adverse effects of protein restriction and resulting in intrauterine growth restriction. By the end of gestation, specifically at 18 days, a significant decrease in fetal weight and cell proliferation was observed, without apparent signs of apoptosis. Autophagy was likely inhibited through the mTOR pathway, potentially conserving resources and protecting functional kidney cells. Thus, in conclusion, gestational protein restriction transiently alters renal growth factor gene expression, initially impacting rapid cell proliferation. Early kidney cell differentiation involves regulators that modulate the dynamics of the autophagy pathway. Overall, these findings suggest that gene regulation, autophagy, and mTOR-mediated mechanisms may substantially reduce nephron numbers following gestational protein restriction beyond the 18th day of gestation.

## ABBREVIATIONS AND ACRONYMS

AngII: angiotensin II.
ANOVA: variance analysis.
AT1: type1 angiotensin receptor
AT2: type 2 angiotensin receptor
BU: Ureteral Bud
BW: body weight.
CAP: metanephric Cap
CEMIB: Multidisciplinary Center for Biological Research in Science in Laboratory Animals.
CEUA: Institutional Ethics Committee approved the experimental design for the use of animals.
CIBIO: Internal Biosafety Commission
CONCEA: Brazilian Council for Animal Experimentation
CREB: cAMP-responsive element-binding
DOHaD: Developmental Origins of Health and Disease
FGF: Fibroblast growth factors
GD: Days of gestation
GDNF: Glial cell line-derived neurotrophic factor
IUGR: intrauterine growth restriction
LP: low-protein intake.
MM: Metanephric Mesenchyme
mTOR: mammalian target of rapamycin.
NP: normal protein intake.
PFA: Paraformaldehyde
Wk: week

## SUPPLEMENTAL FIGURES AND TABLES

**Figure 1S.** Autophagy process at 14 GD (B) and 18 GD © in the NP Group and the LP Offspring. (A). The Lamp1 marker, which indicates lysosomes, emits a blue signal. The combined GFP and RFP fluorescence produces a yellow signal in high pH environments within phagophores and autophagosomes. As maturation proceeds and the vesicles acidify, the EGFP signal diminishes, shifting from yellow to orange to Red. This indicates that EGFP is quenched in mature (M) autophagosomes, leaving only the RFP signal. The autolysosome generates a purple signal (RFP + Lamp1).

**Table 1S.** Composition of standard rodent laboratory diet (standard normal-protein (NP) diet, 17%, and low-protein (LP) diet, 6% (AIN 93G)

**Table 2S.** Antibodies are used at their appropriate concentrations and with the appropriate supplies.

**Table 3S.** Table 3S presents fold changes and p-values for the LP group (experimental group) compared with the NP group (control group) for 84 genes associated with growth factors.

## Ethics approval and consent to participate

The Institutional Ethics Committee (approval #5481-1/2020 CEUA/UNICAMP) authorized the use of transgenic animals, as confirmed by the Internal Biosafety Commission (CIBIO/FCM n° 08/2019). The study utilized C57BL/6J mice and C57BL/6-Tg (CAG-RFP/EGFP/Map1lc3b) 1Hill/J (CAG-RFP-EGFP-LC3). Throughout the investigation, we adhered to the general guidelines established by the Brazilian College of Animal Experimentation.

## Consent for publication

All authors approved for publication.

All primary datasets generated and analyzed in this study are available at http://repositorio.unicamp.br/jspui/handle/REPOSIP/312739?mode=full. Please visit the repository for direct access, review, and download of datasets, in accordance with its terms.

http://repositorio.unicamp.br/jspui/bitstream/REPOSIP/312739/1/_M.pdf

Additional datasets are hosted at http://catalogodeteses.capes.gov.br/catalogo-teses/#!/. Access may require following the repository’s institutional policies. Users should verify the site’s eligibility or coverage requirements.

Data are also available from NCBI’s Gene Expression Omnibus (GEO) via accession GSE188836 (https://www.ncbi.nlm.nih.gov/geo/query/acc.cgi?acc=GSE188836). Please use the GEO portal and abide by NCBI’s access policies.

## Acknowledgments and financial support

MS Folguieri was supported by a CAPES PhD fellowship. Funding was provided by FAPESP (CEPID OCRC, 2013/12486-5 and 2024/06071-1), CNPq (465699/2014-6 and 308043/2022-7), and the Brazilian National Institute of Science and Technology of DOHaD-INCT (409165/2024-0).

## Authors’ contributions

MSF: data curation, investigation, formal analysis, methodology, visualization, writing–original draft; BC: Investigation, methodology, visualization; JARG: conceptualization, funding acquisition, formal analysis, methodology, visualization, writing–review & editing; PAB: conceptualization, funding acquisition, formal analysis, methodology, supervision, visualization, writing–original draft & editing.

## Authors’ information

Marina S. Folguieri, BSC, MS

Department of Internal Medicine, School of Medicine, State University of Campinas, Campinas, SP, Brazil

Phone: +55 19 35217346, Fax: +55 19 3521-8925

Bruno Calsa, BSC, MS

Department of Internal Medicine, School of Medicine, State University of Campinas, Campinas, SP, Brazil

Phone: +55 19 35217346, Fax: +55 19 3521-8925

Patricia Aline Boer, BSC, Ph.D.

Department of Internal Medicine, School of Medicine, State University of Campinas, Campinas, SP, Brazil

Phone: +55 19 35217346, Fax: +55 19 3521-8925

## Acknowledgments

The authors thank Ize Penha Lima for technical assistance.

## Declarations

**Declaration Regarding the Use of Generative AI and AI-Assisted Technologies in Writing - Statement:** The author(s) declare that, during the preparation of this work, no generative AI or AI-assisted technologies were used in the writing process. The author(s) accept full responsibility for the content of this publication.

## REFERENCES

1. Mesquita FF, Gontijo JAR, Boer PA. 2010a. Expression of renin–angiotensin system signalling compounds in maternal protein-restricted rats: effect on renal sodium excretion and blood pressure, Nephrol Dial Transpl, 25(2): 380–388 2010b 10.1093/ndt/gfp505

2. Mesquita FF, Gontijo JAR, Boer PA. 2010b. “Maternal Undernutrition and the Offspring Kidney: From Fetal to Adult Life.” Braz J Med Biol Res, 43 (11): 1010–18. 10.1590/S0100-879X2010007500113.

3. Luyckx, Valerie A., John F. Bertram, Barry M. Brenner, Caroline Fall, Wendy E. Hoy, Susan E. Ozanne, and Bjorn E. Vikse. 2013. “Effect of Fetal and Child Health on Kidney Development and Long-Term Risk of Hypertension and Kidney Disease.” Lancet 382 (9888): 273–83. 10.1016/S0140-6736(13)60311-6.

4. Sene B, Mesquita FF, Santos DC, Carvalho R, Gontijo JAR, Boer PA. Involvement of Renal Corpuscle microRNA Expression on Epithelial-to-Mesenchymal Transition in Maternal Low Protein Diet in Adult Programmed Rats. PLOS ONE, 8(8), 2013, e71310. 10.1371/journal.pone.0071310

5. Short Kieran M, Smyth IM. 2016. “The Contribution of Branching Morphogenesis to Kidney Development and Disease.” Nature Reviews Nephrology 12 (12): 754–67. 10.1038/nrneph.2016.157.

6. Oxburgh Leif. 2018. “Kidney Nephron Determination.” Ann Rev Cell Dev Biol 34: 427–50. 10.1146/annurev-cellbio-100616-060647.

7. Sene B, Scarano WR, Zapparoli A, Gontijo JAR, Boer PA. Impact of gestational low-protein intake on embryonic kidney microRNA expression and in nephron progenitor cells of the male fetus. PLOS ONE, 16(2), 2021, e0246289. 10.1371/journal.pone.0246289

8. Lamana GL, Ferrari ALL, Gontijo JAR, Boer PA. 2021. “Gestational and Breastfeeding Low-Protein Intake on Blood Pressure, Kidney Structure, and Renal Function in Male Rat Offspring in Adulthood.” Front Physiol 12 (April). 10.3389/fphys.2021.658431.

9. Zandi-Nejad K, Luyckx VA, Brenner BM. 2006. “Adult Hypertension and Kidney Disease: The Role of Fetal Programming.” Hypertension 47 (3 II): 502–8. 10.1161/01.HYP.0000198544.09909.1a.

10. Schreuder M, Delemarre-Van De Waal H, Wijk AV. 2006. “Consequences of Intrauterine Growth Restriction for the Kidney.” Kidney Blood Press Res 29 (2): 108–25. 10.1159/000094538.

11. Lucas Alan. 1991. “Programming by Early Nutrition in Man. The Childhood Environment and Adult Disease.” Child Env Adult Dis, 38–55.

12. Lucas Alan. 1998. “Programming by Early Nutrition: An Experimental Approach.” The J Nut, no. April, 407–10.

13. Barker DJ, C. Osmond C, Golding J, Kuh D, Wadsworth ME. 1989. “Growth in *Utero*, Blood Pressure in Childhood and Adult Life, and Mortality from Cardiovascular Disease.” BMJ 298 (6673): 564–67. 10.1136/bmj.298.6673.564.

14. Barker DJP, Osmond C, Winter PD, Margetts B, Simmonds SJ. 1989. “Saturday 9 September 1989 WEIGHT IN INFANCY AND DEATH FROM ISCHAEMIC HEART DISEASE.” Lancet 334 (8663): 577–80. 10.1016/S0140-6736(89)90710-1.

15. Zeman FJ. 1967. “Effect of the Young Rat of Maternal Protein Restriction.” J Nut 93 (2): 167–73. 10.1093/jn/93.2.167.

16. Langley-Evans SC, Gardner DS, Jackson AA. 1996. “Association of Disproportionate Growth of Fetal Rats in Late Gestation with Raised Systolic Blood Pressure in Later Life.” Reproduction 106 (2): 307–12. 10.1530/jrf.0.1060307.

17. Langley-Evans SC, Simon Welham SJM, Jackson AA. 1999. “Fetal Exposure to a Maternal Low Protein Diet Impairs Nephrogenesis and Promotes Hypertension in the Rat.” Life Sci 64 (11): 965–74. 10.1016/S0024-3205(99)00022-3.

18. Merlet-Bénichou C, Gilbert T, Muffat-Joly M, Lelièvre-Pégorier M, Leroy B. 1994. “Intrauterine Growth Retardation Leads to a Permanent Nephron Deficit in the Rat.” Ped Nephrol 8 (2): 175–80. 10.1007/BF00865473.

19. Kasture V, Sahay A, Joshi S. 2021. *Cell Death Mechanisms and Their Roles in Pregnancy-Related Disorders*. Adv Prot Chem Struc Biol. 1st ed. Vol. 126. Elsevier Inc. 10.1016/bs.apcsb.2021.01.006.

20. Westerterp-Plantenga MS, Nieuwenhuizen A, Tomé D, Soenen S, Westerterp KR. 2009. “Dietary Protein, Weight Loss, and Weight Maintenance.” Ann Rev Nut 29 (2009): 21–41. 10.1146/annurev-nutr-080508-141056.

21. Westerterp-Plantenga MS, Nieuwenhuizen A, Soenen S, Westerterp KR, Tome D, Marion J, et al. 2009. “Brain Responses to High-Protein Diets.” Ann Rev Nut 92 (2009): S27–30. 10.1146/annurev-nutr-080508-141056.

22. Davidenko O, Darcel N, Fromentin G, Tomé D. 2013. “Control of Protein and Energy Intake - Brain Mechanisms.” Eur J Clin Nutr 67 (5): 455–61. 10.1038/ejcn.2013.73.

23. Tomé D. 2004. “Protein, Amino Acids and the Control of Food Intake.” Brit J Nutr 92 (S1): S27–30. 10.1079/bjn20041138.

24. Desai M, Crowther NJ, Lucas A, Hales CN. 1996. “Organ-Selective Growth in the Offspring of Protein-Restricted Mothers.” Brit J Nutr 76 (4): 591–603. 10.1079/bjn19960065.

25. Passos MCF, Ramos CF, Moura EG. 2000. “SHORT AND LONG TERM EFFECTS OF MALNUTRITION IN RATS DURING LACTATION ON THE BODY WEIGHT OF OFFSPRING M.” Nutr Res 20 (I): 1603–1612.

26. Colombo JP, Cervantes H, Kokorovic M, Pfister U, Perritaz R. 1992. “Effect of Different Protein Diets on the Distribution of Amino Acids in Plasma, Liver and Brain in the Rat.” Ann Nutr Metab 4 (1): 23–33.

27. Bell RR, McGill TJ, Digby PW, Bennett SA. 1984. “Effects of Dietary Protein and Exercise on Brown Adipose Tissue and Energy Balance in Experimental Animals.” Journal of Nutrition 114 (10): 1900–1908. 10.1093/jn/114.10.1900.

28. Toyomizu M, Matsukubo M, Hayashi K, Tomita Y. 1991. “Response Surface Analyses of the Effects of Dietary Fat on Feeding and Growth Pattern in Mice from Weaning to Maturity.” Animal Prod 52 (1): 207–14. 10.1017/S0003356100005857.

30. Aparecida de França S, dos Santos MP, Maria Garófalo MAR, Navegantes LC, Kettelhut IC, Lopes CF, Kawashita NH. 2009. “Low Protein Diet Changes the Energetic Balance and Sympathetic Activity in Brown Adipose Tissue of Growing Rats.” Nutrition 25 (11–12): 1186–92. 10.1016/j.nut.2009.03.011.

31. Corstius HB, Zimanyi MA, Maka N, Herath T, Thomas W, Der Laarse AV, Wreford NG, Black MJ. 2005. “Effect of Intrauterine Growth Restriction on the Number of Cardiomyocytes in Rat Hearts.” Ped Res 57 (6): 796–800. 10.1203/01.PDR.0000157726.65492.CD.

32. Rees WD, Hay SM, Buchan V, Antipatis C, Palmer RM. 1999. “The Effects of Maternal Protein Restriction on the Growth of the Rat Fetus and Its Amino Acid Supply.” Brit J Nut 81 (3): 243–50. 10.1017/s0007114599000446.

33. Fowden AL, Sferruzzi-Perri AN, Coan PM, Constancia M, Burton GJ. 2009. “Placental Efficiency and Adaptation: Endocrine Regulation. J Physiol, 587: 3459–72. 10.1113/jphysiol.2009.173013.

34. Hoppe CC, Evans RG, Bertram JF, Moritz KM. 2007. “Effects of Dietary Protein Restriction on Nephron Number in the Mouse.” Am J Physiol - Regulatory Integrative and Comparative Physiology 292 (5): 1768–75. 10.1152/ajpregu.00442.2006.

35. Woods, L, Ingelfinger, J.R., Nyengaard, J.R., Rasch, R. 2001. “Maternal Protein Restriction Suppresses the Newborn Renin-Angiotensin System and Programs Adult Hypertension in Rats.” Ped Res 49 (4): 460–67. 10.1203/00006450-200104000-00005.

36. Woods L, Ingelfinger JR, Nyengaard JR, Rasch R, Hoppe CC, Evans RG, Bertram JF, et al. 2022. “Effects of Dietary Protein Restriction on Nephron Number in the Mouse.” Am J Physiol - Regulatory Integrative and Comparative Physiology 49 (20): 460–67. 10.1152/ajpregu.00442.2006.

37. Pezzotta A, Perico L, Morigi M, Corna D, Locatelli M, Zoja C, Benigni A, Remuzzi G, Imberti B. 2022. “Low Nephron Number Induced by Maternal Protein Restriction Is Prevented by Nicotinamide Riboside Supplementation Depending on Sirtuin 3 Activation.” Cells 11 (20). 10.3390/cells11203316.

38. Reena S, Watanabe T, Costantini F. 2005. “The Role of GDNF/Ret Signaling in Ureteric Bud Cell Fate and Branching Morphogenesis.” Devel Cell 8 (1): 65–74. 10.1016/j.devcel.2004.11.008.

39. Costantini F. 2010. “GDNF/Ret Signaling and Renal Branching Morphogenesis.” Organogenesis 6 (4): 252–62. 10.4161/org.6.4.12680.

40. Costantini F, Shakya R. 2006. “GDNF/Ret Signaling and the Development of the Kidney.” BioEssays 28 (2): 117–27. 10.1002/bies.20357.

41. Godin RE, Robertson EJ, Dudley AT. 1999. “Role of BMP Family Members during Kidney Development.” Int J Dev Biol 43 (5): 405–11.

42. Ok-Hee C, Chang-Ho Song, Sung-Kwang Park, Won Kim, and Eui-Sic Cho. 2013. “Molecular Regulation of Kidney Development.” Anat Cell Biol 46 (1): 19. 10.5115/acb.2013.46.1.19.

43. Mayumi T, Asada M, Asada N, Nakamura J, Oguchi A, Atsuko Y. Higashi, Endo S, et al. 2013. “Bmp7 Maintains Undifferentiated Kidney Progenitor Population and Determines Nephron Numbers at Birth.” PLoS ONE 8 (8): 1–9. 10.1371/journal.pone.0073554.

44. Khoshdel RN, Aghdami N, Moghadasali R. 2020. “Cellular and Molecular Mechanisms of Kidney Development: From the Embryo to the Kidney Organoid.” Front Cell Develop Biol 8 (March): 1–16. 10.3389/fcell.2020.00183.

45. Egan RJ, 2009. “NIH Public Access.” Behavioral Brain Res 205 (9): 38–44. 10.1007/s00467-011-1819-8.

46. Carev D, Saraga M, Saraga-Babic M. 2008. “Expression of Intermediate Filaments, EGF and TGF-α in Early Human Kidney Development.” J Mol Histol 39 (2): 227–35. 10.1007/s10735-007-9157-7.

47. Walker KA, Sims-Lucas S, Bates CM. 2016. “Fibroblast Growth Factor Receptor Signaling in Kidney and Lower Urinary Tract Development.” Pediatr Nephrol. 31 (6): 885–895. 10.1007/s00467-015-3151-1.Fibroblast.

48. Xie Y, Su N, Yang J, Tan Q, Huang S, Jin M, Ni Z, et al. 2020. “FGF/FGFR Signaling in Health and Disease.” Signal Trans Targ Ther 5 (1). 10.1038/s41392-020-00222-7.

49. Peranton AO, Timofeeva O, Naillat F, Richman C, Pajni-Underwood S, Wilson C, Vainio S, Dove LF, Lewandoski M. 2005. “Inactivation of FGF8 in Early Mesoderm Reveals an Essential Role in Kidney Development.” Development 132 (17): 3859–71. 10.1242/dev.01945.

50. Cancilla B, Ford-Perriss MD, Bertram JF. 1999. “Expression and Localization of Fibroblast Growth Factors and Fibroblast Growth Factor Receptors in the Developing Rat Kidney.” Kidney Intern 56 (6): 2025–39. 10.1046/j.1523-1755.1999.00781.x.

51. Grieshammer U, Cebrián C, Ilagan R, Meyers E, Herzlinger D, Martin GR. 2005. “FGF8 Is Required for Cell Survival at Distinct Stages of Nephrogenesis and for Regulation of Gene Expression in Nascent Nephrons.” Development 132 (17): 3847–57. 10.1242/dev.01944.

52. Huber L. Julie, Hempstead B, Donovan MJ. 1996. “Neurotrophin and Neurotrophin Receptors in Human Petal Kidney.” Devel Biol 179 (2): 369–81. 10.1006/dbio.1996.0268.

53. Abe T, Fleming PA, Masuya M, Minamiguchi H, Ebihara Y, Drake CJ, Ogawa M. 2005. “Granulocyte/Macrophage Origin of Glomerular Mesangial Cells.” Int J Hematol 82 (2): 115–18. 10.1532/IJH97.05018.

54. Masuya M, Drake CJ, Fleming PA, Reilly CM, Zeng H, Hill WD, Martin-Studdard A, Hess DC, Makio Ogawa M. 2003. “Hematopoietic Origin of Glomerular Mesangial Cells.” Blood 101 (6): 2215–18. 10.1182/blood-2002-04-1076.

55. Sauter, Kristin A, Clare Pridans, Anuj Sehgal, Yi Ting Tsai, Barry M Bradford, Sobia Raza, Lindsey Moffat, et al. 2014. “Pleiotropic Effects of Extended Blockade of CSF1R Signaling in Adult Mice.” J Leuk Biol 96 (2): 265–74. 10.1189/jlb.2a0114-006r.

56. Rae F, Woods K, Sasmono T, Campanale N, Taylor D, Ovchinnikov DA, Grimmond SM, Hume DA, Ricardo SD, Little MH. 2007. “Characterisation and Trophic Functions of Murine Embryonic Macrophages Based upon the Use of a Csf1r-EGFP Transgene Reporter.” Devel Biol 308 (1): 232–46. 10.1016/j.ydbio.2007.05.027.

57. Alikhan, MA, Jones, C.V., Williams, T.M., Beckhouse, A.G., Fletcher AL, Kett MM, Sakkal, S., et al. 2011. “Colony-Stimulating Factor-1 Promotes Kidney Growth and Repair via Alteration of Macrophage Responses.” Am J Pathol 179 (3): 1243–56. 10.1016/j.ajpath.2011.05.037.

58. Munro DA, Wineberg Y, Tarnick J, Vink CS, Li Z, Pridans C, Dzierzak E, Kalisky T, Hohenstein P, Davies JA. 2019. “Macrophages Restrict the Nephrogenic Field and Promote Endothelial Connections during Kidney Development.” ELife 8: 1–27. 10.7554/eLife.43271.

59. Sutherland MR., Black MJ. 2023. “The Impact of Intrauterine Growth Restriction and Prematurity on Nephron Endowment.” Nat Rev Nephrol 19 (4): 218–28. 10.1038/s41581-022-00668-8.

60. Wang C, Zhang R, Zhou L, He J, Huang Q, Siyal FA, Zhang L, Zhong X, Wang T. 2017. “Intrauterine Growth Retardation Promotes Fetal Intestinal Autophagy in Rats via the Mechanistic Target of Rapamycin Pathway.” J Reprod Develop 63 (6): 547–54. 10.1262/jrd.2017-050.

61. Stewart T, Kallash M, Vehaskari VM, Hodgeson SM, Aviles DH. 2019. “Increased Autophagy and Apoptosis in the Kidneys of Intrauterine Growth Restricted Rats.” Fetal Ped Pathol 38 (3): 185–94. 10.1080/15513815.2018.1564160.

62. Eskelinen EL. 2005. “Maturation of Autophagic Vacuoles in Mammalian Cells.” Autophagy 1 (1): 1–10. 10.4161/auto.1.1.1270.

63. Dunn WA. 1990. “Studies on the Mechanisms of Autophagy: Maturation of the Autophagic Vacuole.” J Cell Biol 110 (6): 1935–45.

64. Li L, Wang ZV, Hill JA, Lin F. 2014. “New Autophagy Reporter Mice Reveal Dynamics of Proximal Tubular Autophagy.” J Am Soc Nephrol 25 (2): 305–15. 10.1681/ASN.2013040374.

65. Liang X, Yu C, Tian Y, Xiang X, Luo Y. 2023. “Inhibition of STX17–SNAP29–VAMP8 Complex Formation by Costunolide Sensitizes Ovarian Cancer Cells to Cisplatin via the AMPK/MTOR Signaling Pathway.” Biochem Pharmacol 212 (June):115549. 10.1016/J.BCP.2023.115549.

66. Ratto E, Chowdhury SR, Siefert NS, Schneider M, Wittmann M, Helm D, Palm W. 2022. “Direct Control of Lysosomal Catabolic Activity by MTORC1 through Regulation of V-ATPase Assembly.” Nat Comm 13 (1). 10.1038/s41467-022-32515-6.

67. Pattingre S, Tassa A, Qu X, Garuti R, Xiao HL, Mizushima N, Packer M, Schneider MD, Levine B. 2005a. “Bcl-2 Antiapoptotic Proteins Inhibit Beclin 1-Dependent Autophagy.” Cell 122 (6): 927–39. 10.1016/j.cell.2005.07.002.

68. Pattingre S, Tassa A, Qu X, Garuti R, Xiao HL, Mizushima N, Packer M, Schneider MD, Levine B. 2005b. “Bcl-2 Antiapoptotic Proteins Inhibit Beclin 1-Dependent Autophagy.” Cell 122 (6): 927–39. 10.1016/j.cell.2005.07.002.

69. Marquez RT, Xu L. 2012. “Bcl-2: Beclin 1 Complex: Multiple Mechanisms Regulating Autophagy/Apoptosis Toggle Switch.” Am J Cancer Res 2 (2):214–221. http://www.ncbi.nlm.nih.gov/pubmed/22485198 http://www.pubmedcentral.nih.gov/articlerender.fcgi?artid=PMC3304572.

70. Decuypere JP, Parys JB, Bultynck G. 2012. “Regulation of the Autophagic Bcl-2/Beclin 1 Interaction.” Cells. MDPI. 10.3390/cells1030284.

71. Gomes JS, Sene LB, Lamana GL, Boer PA, Gontijo JAR. Impact of maternal protein restriction on Hypoxia-Inducible Factor (HIF) expression in male fetal kidney development. PLOS ONE, 18(5), e0266293. 10.1371/journal.pone.0266293

72. Tsuji K, Kitamura S, Makino H. 2014. “Hypoxia-Inducible Factor 1α Regulates Branching Morphogenesis during Kidney Development.” Biochem Biophys Res Commun 447(1): 108–14. 10.1016/j.bbrc.2014.03.111.

73. Gunaratnam L, Bonventre JV. 2009. “HIF in Kidney Disease and Development.” J Am Soc Nephrol 20 (9): 1877–87. 10.1681/ASN.2008070804.

74. Schley G, Scholz H, Kraus A, Hackenbeck T, Klanke B, Willam C, Wiesener MS, et al. 2015. “Hypoxia Inhibits Nephrogenesis through Paracrine Vegfa despite the Ability to Enhance Tubulogenesis.” Kidney Int 88 (6): 1283–92. 10.1038/ki.2015.214.

75. Gille J, Swerlick RA, Caughman SW. 1997. “Transforming Growth Factor-α-Induced Transcriptional Activation of the Vascular Permeability Factor (VPF/VEGF) Gene Requires AP-2-Dependent DNA Binding and Transactivation Also Known as Vascular Endothelial Growth Factor (VEGF).” EMBO J. Vol. 16.

76. Ohki Y, Heissig B, Sato Y, Akiyama H, Zhu Z, Hicklin DJ, Shimada K, et al. 2005. “Granulocyte Colony-stimulating Factor Promotes Neovascularization by Releasing Vascular Endothelial Growth Factor from Neutrophils.” FASEB J 19 (14): 2005–7. 10.1096/fj.04-3496fje.

